# RoBep: A Region-Oriented Deep Learning Model for B-Cell Epitope Prediction

**DOI:** 10.1101/2025.11.22.689912

**Authors:** Yitao Xu, Guanyun Wei, Jingying Zhou, Yuanhua Huang, Weichua Yu, Zhixiang Lin, Ran Liu, Xiaodan Fan

## Abstract

**Motivation:** Accurate in silico identification of B-cell epitope residues is crucial for antibody design and structure-guided vaccine development. Although recent protein language models and structure-aware methods can capture spatial information of tertiary structure when generating residue embeddings, most existing epitope predictors use these embeddings to perform classification for individual residues one by one, without enforcing spatial continuity for reported epitope residues. Such methods often result in biologically implausible predictions because B-cell epitope residues always cluster together on the antigen surface.

**Results:** We present RoBep, a region-oriented B-cell epitope predictor that explicitly models the spatial clustering of epitope residues. RoBep introduces a novel region constraint mechanism and combines the advanced protein language model ESM-Cambrian with an equivariant graph neural network. Our method outperforms existing structure-based methods on the benchmark dataset, demonstrating improvements of 26%, 45%, 13%, and 43% in F1, MCC, AUPR, and AUROC_0.1_, respectively. In addition to residue-level predictions, RoBep can also provide antibody-antigen binding regions. Importantly, the predicted epitope residues are ensured to be spatially compact, enhancing biological plausibility and practical relevance for immunotherapeutic design.

**Availability:** A user-friendly website for using RoBep is provided at https://huggingface.co/spaces/NielTT/RoBep. All datasets, source code used in this work, and implementation instructions of the website are publicly available at https://github.com/YitaoXU/RoBep.

## 1 Introduction

B cells are central to the humoral immune response by producing antibodies that bind to and neutralize potentially harmful antigens [11]. The specific residues of an antigen recognized by an antibody are termed B-cell epitopes (BCEs). BCEs can be categorized as either linear, contiguous in the primary sequence or conformational, composed of discontinuous residues brought into proximity by the protein’s three-dimensional structure, with the latter accounting for over 90% of all BCEs [1]. Accurate identification of these epitopes is essential for understanding immune mechanisms [10], as well as advancing applications such as mRNA vaccine development [30] and antibody design [4]. For example, the de novo antibody design model Chai-2 requires antigen target residues as input, underscoring the importance of precise conformational BCE prediction. Therefore, reliable localization of such epitopes is crucial for both basic immunology research and relevant applications.

Current experimental methods, such as X-ray crystallography and nuclear magnetic resonance, offer reliable epitope mapping but are costly and time-consuming, limiting scalability [13]. This has spurred the development of in silico approaches for B-cell epitope prediction. Early in silico methods typically relied on handcrafted features derived from structural and physicochemical properties, fed into machine learning models like support vector machines. For example, the DiscoTope series [24] used features such as solvent accessibility and contact number, while the SEPPA series [40] added biological factors like glycosylation. However, their overall performance remained unsatisfactory [7], likely due to data scarcity and the complexity of immune recognition, which handcrafted descriptors fail to effectively capture.

The recent emergence of large protein language models (PLMs) [17, 28, 35], based on the transformer architecture [37], has markedly enriched the sequence- and structure-level representations available for downstream tasks, including epitope prediction. Leveraging these models, many recent BCE prediction approaches have achieved substantial advances. Several methods adopted the sequence-based PLM ESM-2 [28] to generate residue-level embeddings, which were then passed to classifiers for epitope prediction [8, 19]. GraphBepi [39] further incorporated structural context through graph neural networks, enhancing predictive accuracy by modeling residue–residue spatial relationships. Meanwhile, methods such as SEMA-2.0 [20] and DiscoTope-3.0 [16], both of which leverage structure-aware PLMs, have demonstrated impressive performance, reaffirming the importance of structural information in predicting conformational BCEs. Nonetheless, earlier work remained largely sequence-centric and focused on linear epitopes, since antigen structures were often unavailable at prediction time in real applications and experimentally determined antigen–antibody complexes providing true conformational BCE for training were scarce.

These limitations are being alleviated by advances on two fronts: AlphaFold [21], which enables accurate structure prediction directly from sequence and thus makes structure-based models widely accessible at inference time, and single-particle cryo-EM [6], which has accelerated the accumulation of experimentally resolved antigen-antibody complexes to expand structure-mapped epitopes. Consistent with these trends, recent studies [16, 39] report comparable performance for conformational BCE prediction when using either experimental or AlphaFold-predicted structures, further motivating structure-guided predictors.

Despite these advances, most existing methods incorporate spatial patterns by extracting informative features for each residue, but feed all features equally into the classifier during the final prediction step. This overlooks a key biological property of BCEs, their natural tendency to form spatially clustered surface regions rather than being dispersed across the whole antigen surface. Reis et al. [31] observed that, in most antigens, the epitope surface typically forms a single connected patch, which can often be covered by an elliptical plane [25]. These findings suggest that spatial compactness is a defining characteristic of BCEs. Ignoring this constraint often leads to fragmented and unrealistic epitope predictions, limiting the utility of current tools in downstream applications such as antibody design. Motivated by this limitation, we aim to develop a biologically consistent epitope predictor that enforces spatial coherence and harnesses modern deep learning techniques to enhance predictive accuracy and structural awareness.

In this work, we present a **R**egion-**o**riented **B**-cell **e**pitope **p**redictor (RoBep), which introduces a novel region constraint mechanism to promote spatially coherent predictions. RoBep integrates multiple deep learning components, including the sequence-based PLM ESM-Cambrian (ESM-C) [15] and enhanced Equivariant Graph Neural Networks (EGNN) [33], to effectively capture both sequence and structural features of antigens. Beyond providing residue-level epitope predictions, RoBep is also capable of identifying high-likelihood antibody–antigen interface regions, offering richer biological insights. Moreover, we demonstrate through comprehensive evaluations that RoBep exhibits high specificity on antigenic proteins, further supporting its practical relevance. Our model not only ensures more realistic spatial distributions of predicted epitope residues but also achieves the best performance on curated benchmark datasets, underscoring its utility for both accurate prediction and downstream immunological applications.

## 2 Materials and Methods

### 2.1 Dataset

We constructed a high-quality antigen dataset for structure-based B-cell epitope prediction following the strategy of Shashkova et al. [34]. All protein complexes from the PDB [2] with a resolution smaller than 4.0 Å (as of November 4, 2024) were retrieved. Antibody-antigen (Ab-Ag) complexes containing heavy or light chains were identified using ANARCI [12], and only those with both chains present were retained. In each complex, the remaining protein chains were designated as candidate antigen chains and processed independently, including multi-chain antigens. All chains with sequence lengths outside 12–2046 residues were removed. For each retained antigen, a residue was labeled as an epitope if any of its heavy atoms were within 4.0 Å of a heavy atom from an antibody, as defined by Kringelum et al. [24]. In addition, antigens with fewer than five epitope residues were excluded. To reduce redundancy, antigen sequences were then clustered at 95% identity using CD-HIT [27], and within each cluster, sequences were aligned using MAFFT [23]. The longest sequence in each cluster was selected as the representative, and epitope annotations from other members were propagated based on aligned positions. This merging step prevents highly similar antigen sequences from carrying discrepant epitope labels into the training data, thereby avoiding harmful supervision conflicts. The curated dataset consisted of 756 antigens comprising 176,846 residues, of which 11,848 (6.7%) were labeled as epitopes, including both linear and conformational BCEs. Finally, the dataset was split into training and test sets by re-running CD-HIT on the non-redundant set at a 70% identity threshold and assigning entire clusters to the two splits to approximate an 8:2 ratio by sequence count, with all member sequences of each assigned cluster included in that split. We additionally double-checked the split with CD-HIT-2D at 70% identity to confirm that no cross-split pairs reach ≥ 70% identity. This procedure yielded 612 antigens for training (140,106 residues, 6.6% epitopes) and 144 for testing (36,740 residues, 7.0% epitopes), ensuring that no antigen in the training set shares ≥ 70% sequence identity with any antigen in the test set and thus avoiding information leakage.

Meanwhile, for the protein-level antigenicity evaluation in Section 3.5, we constructed an auxiliary antigenicity dataset comprising 144 antigens (positives) from the test set and 85 non-structural proteins from viruses (negatives). This choice follows the fact that non-structural proteins (nsps) generally exhibit low antigenicity and are rarely recognized by antibodies [18]. This dataset is used solely to assess RoBep’s ability to distinguish antigenic proteins from weakly antigenic nsps. Further details are provided in Supplementary Materials S1.

### 2.2 RoBep Framework

The overall architecture of RoBep is illustrated in Figure 1. Given an antigen structure, RoBep first enumerates candidate antibody–antigen binding regions using a rolling-sphere strategy. Each region is modeled as a residue-level graph, with node features combining ESM-C embeddings and structural descriptors, and edge features capturing spatial and sequential relationships. Two layers of E(n)-equivariant Graph Neural Network (EGNN) encode these graphs to produce structure-aware residue representations. Region-level scores are then computed via attention pooling, and the top-ranked regions are selected. Residue-level epitope prediction is performed by integrating residue and region embeddings through a multilayer perceptron module (MLP), with the final probability for each residue obtained by averaging its predictions across all regions in which it appears. All predicted epitope residues lie entirely within the selected regions, a design referred to as the region constraint mechanism, which encourages spatial coherence. Detailed descriptions of each component are provided in the following subsections.

**Figure 1.**
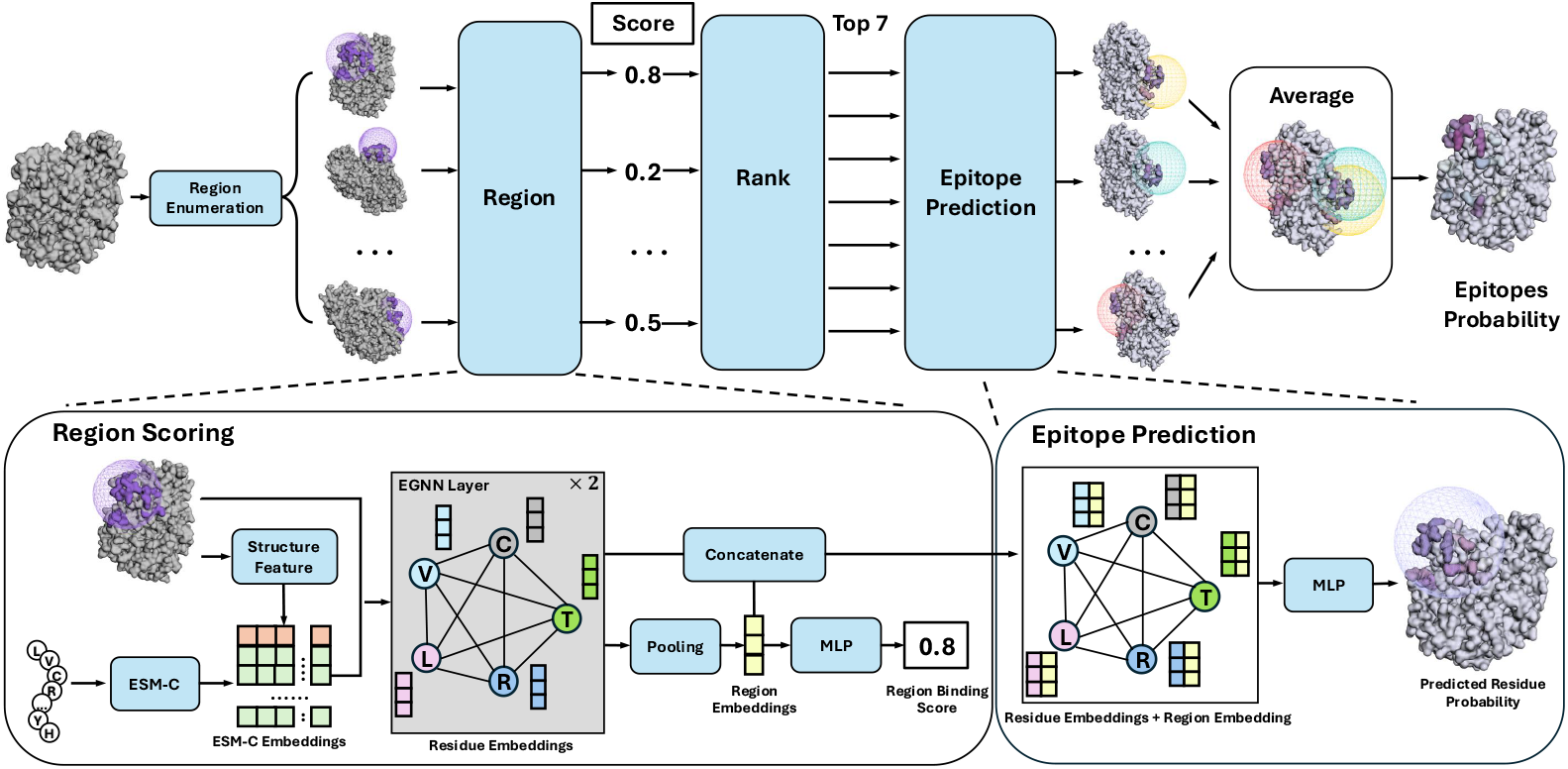
Overview of the RoBep framework for B-cell epitope prediction on a single antigen. The model identifies candidate surface regions using a rolling-sphere strategy, encodes each region as a residue-level graph with both sequence-based and structure-based features, and applies an E(n)-equivariant GNN to extract residue representations. High-confidence regions are scored and selected for downstream residue-level probability prediction, which integrates both local and regional information. The final epitope probability for each residue is obtained by averaging its predictions across all selected regions.

#### 2.2.1 Region Enumeration

To support the region constraint mechanism, we first enumerate candidate surface regions that may serve as potential Ab-Ag binding regions. We employ a sphere-based scanning strategy ^1^: the C*α* atoms of all solvent-accessible residues^2^ are selected as sphere centers, and spheres with a fixed radius of *r* = 18 Å are constructed around each residue.

Each candidate region is defined as the set of residues whose C*α* atoms fall within the intersection of the sphere and the antigen surface. The radius is chosen based on ag-ab interface statistics (mean area 1068 314 ± Å^2^ [31]), ensuring adequate coverage for most epitopes. All these candidate regions are subsequently used for scoring and residue-level prediction. To enforce spatial coherence, final residue predictions are constrained to lie within selected high-scoring regions, forming the basis of our region constraint mechanism.

#### 2.2.2 Geometric Graph Encoding

Each candidate region is represented as a fully connected residue-level graph *G* = ( 𝒱, ℰ ), where nodes correspond to residues and edges connect all residue pairs within the region. Node features are composed of (i) projected ESM-C embeddings, derived from the 2560-dimensional pretrained output of ESM-C [15], mapped to the hidden dimension via an MLP, and (ii) structural descriptors, including RSA and backbone dihedral angles. Edge features include (i) spatial distances, transformed from the Euclidean distance between *C*_*α*_ atoms using 16 Gaussian radial basis functions, and (ii) sequence offsets, encoded as 16-dimensional sinusoidal positional embeddings following Vaswani et al. [37]. For a given graph, we denote the initial node feature of residue *i* as 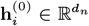, and the edge feature between residue *i* and *j* as 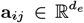. Details of node and edge feature construction are provided in the Supplementary Materials S2.

For simplicity, we use *ϕ*( · ) to denote an MLP, where each instance (e.g., *ϕ*_ESM_, *ϕ*_*e*_, *ϕ*_*h*_) has its own learnable parameters. Unless otherwise specified, MLPs are composed of linear layers with activation functions, followed by dropout and normalization for regularization and stability.

To capture the geometric context of each region, RoBep applies *L* layers of EGNNs [33], enhanced with architectural improvements inspired by Luo et al. [29]. At each layer *l*, each residue *i* is associated with a hidden representation 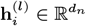 and a 3D coordinate 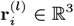. Let 𝒩 (*i*) denote the set of neighboring residues within the region (all other nodes). For every residue pair (*i, j*), the message is computed based on the current node features, coordinates, and edge attributes:

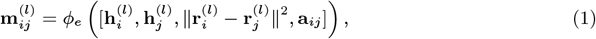

where *ϕ*_*e*_ is an MLP. The 3D coordinate of residue *i* is then updated via a weighted sum of relative displacements:

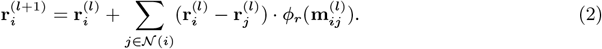

To update node features, the aggregated messages are fused with the current representation through two MLPs:

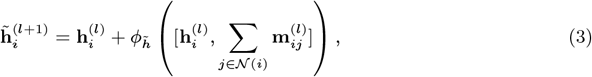

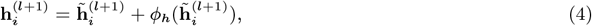

where 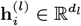 is the node embeddings of residue *i* at layer *l* and all *ϕ*( · ) denote MLPs with separate parameters. The node embeddings 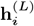 produced by the last EGNN layer are used for both region- and residue-level prediction tasks in subsequent steps.

#### 2.2.3 Region Scoring and Ranking

To predict the proportion of epitopes 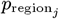 in each candidate region *j*, we apply attention-based pooling over the set of residue embeddings **h**_*i*_ in this region to obtain a global region representation **h**^reg^ ∈ ℝ^256^:

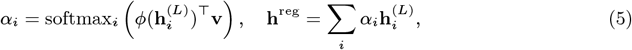

where *α*_*i*_ ∈ ℝ are attention weights and **v** is a learnable vector. The pooled vector is then transformed into a region-level proportion:

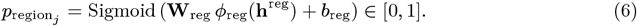

All candidate regions are ranked by 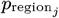, and the top-7 regions, region_(1)_, …, region_(7)_, are retained for residue-level prediction. This step enforces the Region Constraint by restricting residue predictions to these selected regions, ensuring spatial coherence and avoiding scattered outputs across the antigen surface. The selection of the value of *k* (number of top-ranked regions) is detailed in Supplementary Material S3.

#### 2.2.4 Epitope Prediction

For each residue *i* within a selected region *j*, we concatenate its embedding 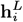 with the region-level representation 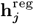, and compute the epitope probability as:

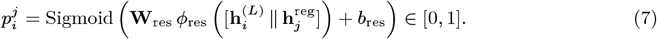

This design allows residue-level predictions to incorporate both local features and global region context. For residues appearing in multiple selected regions, we average their predictions:

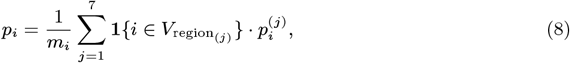

where *m*_*i*_ is the number of selected regions that include residue *i* and 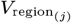 is the set of residues in region_(*j*)_. Residues not included in any selected region are assigned zero.

### 2.3 Objective Function

To effectively train both region- and residue-level predictions, we adopt a joint loss function with dynamic weights:

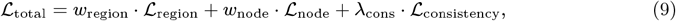

where *w*_region_ and *w*_node_ are loss weights dynamically adjusted via GradNorm [5], which is detailed in Supplementary Materials S4. *λ*_cons_ controls the strength of the consistency term. Each loss component is defined below.

#### Region-level loss

For each candidate region *j*, we supervise the predicted proportion 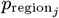 of epitopes in the region using a target-weighted mean squared error (MSE):

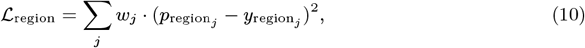

where 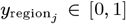 is the fraction of true epitope residues in region *j*, and *w*_*j*_ is a weight that increases with 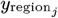 encouraging accurate prediction of highly enriched regions.

#### Residue-level loss

For each residue *i* in region *j*, the predicted epitope probability 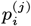 is supervised with focal loss:

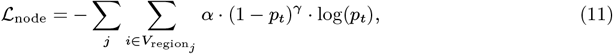

where 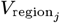 denotes the set of residues in region *j*, and 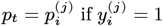, otherwise 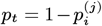. This formulation encourages the model to focus on sparse and difficult-to-identify epitopes.

#### Cross-level consistency

To align region- and residue-level predictions, we include a consistency term:

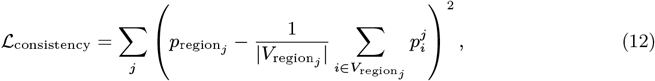

which aligns each region’s proportion with its residue-level scores, encouraging mutual consistency. This multi-level loss provides complementary supervision across levels, and reinforces region–residue consistency, improving prediction accuracy.

### 2.4 Antigenicity Prediction

For any given protein *k*, we define its antigenicity score as the average predicted epitope probability across residues within its top-7 predicted regions. Formally, the score is computed as:

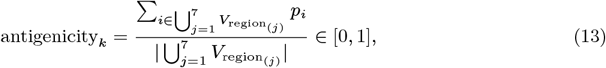

where 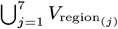 denotes the union of residues in the top-7 region_(*j*)_ selected in Section 2.2.1 and *p*_*i*_ denotes the predicted probability of residue *i* being an epitope in Section 2.2.4. This score predicts the antigenicity of a given protein.

### 2.5 Evaluation Metrics and Experiment Setting

To comprehensively evaluate our method against baselines, we report five metrics: F1, Matthews correlation coefficient (MCC), area under the precision–recall curve (AUPR), area under the ROC curve at low false-positive rates (AUROC_0.1_) [32], and an epitope–overlap metric called the antigen-level intersection over union (AgIoU) [38]. All these metrics can capture model performance well in scenarios with significant class imbalance, given that epitope residues constitute only 6.7% of the entire dataset.

To further evaluate the spatial coherence of predicted epitopes, we calculate the clustered mean pairwise distance (cMPD) between C_*α*_ atoms of predicted residues for each antigen, a metric modified from Colavin et al. [9] that provides a fair and accurate assessment of residues compactness, including multi-region cases. Lower cMPD values indicate tighter clustering of predicted residues, reflecting higher biologically plausible prediction results. We also compare the predicted cluster count (NoC), the number of distinct epitope regions identified by each model, to the ground-truth NoC to assess regional correspondence. Details of metrics are provided in Supplementary Materials S5.

Models were trained with a batch size of 64 for up to 50 epochs, using early stopping with a patience of 10. We used the AdamW optimizer with a maximum learning rate of 5 × 10^−5^, a 10% linear warm-up, and cosine annealing with restarts. All experiments were run on two NVIDIA Quadro GV100 GPUs. More training implementation details are provided in Supplementary Materials S6.

## 3 Results

### 3.1 Benchmark Comparison

We compared our method, RoBep, with popular state-of-the-art methods for structure-based prediction of conformational B-cell epitopes on the test set. These methods leverage structural information through various strategies, including handcrafted physicochemical features (e.g., Seppa-3.0), structure-based protein language models (PLMs; e.g., SEMA-2.0, DiscoTope-3.0), and GNN (e.g., GraphiBepi). For CALIBER, we used the best-performing reproducible configuration (ESM-2 + BiLSTM) reported in the original paper of Israeli and Louzoun [19]. To ensure a fair comparison, we select the decision threshold for each method by sweeping *τ* ∈ [0, 1] (step 0.05) to maximize its F1, and then fix that *τ* for all thresholded metrics.

Since some existing methods, such as GraphiBepi and CALIBER, adopt earlier-generation sequence-level PLMs like ESM-2 as encoder, we also evaluated our model using the same encoder to ensure a fair comparison. As summarized in Table 1, our model still consistently outperformed all baselines under this setting, with improvements of 10.5% in AUPRC, 15.3% in F1 score, and 18.1% in AgIoU and 27.3% in AUROC_0.1_ over the strongest competitor (DiscoTope-3.0). Figure 2 further illustrates this advantage by showing that our method achieves the highest precision across almost the entire recall range. These results highlight the effectiveness of our region-constrained design, which not only enhances biological plausibility but also enhances epitope localization accuracy.

**Table 1:** Performance comparison of our model with state-of-the-art methods with variant PLMs on the independent test dataset.

| Model | PLM | F1 | MCC | AgIoU | AUPR | AUROC<sub>0.1</sub> | Threshold |
| --- | --- | --- | --- | --- | --- | --- | --- |
| Seppa-3.0 | - | 0.194 | 0.131 | 0.107 | 0.118 | 0.125 | 0.15 |
| SEMA-2.0 | SaProt | 0.284 | 0.196 | 0.166 | 0.236 | 0.201 | 0.75 |
| GraphiBepi | ESM-2 | 0.276 | 0.189 | 0.156 | 0.215 | 0.172 | 0.15 |
| DiscoTope-3.0 | ESM-IF | 0.300 | 0.234 | 0.177 | 0.237 | 0.231 | 0.30 |
| CALIBER | ESM-2 | 0.241 | 0.164 | 0.137 | 0.158 | 0.156 | 0.10 |
| RoBep | ESM-2 | 0.346 | 0.295 | 0.209 | 0.262 | 0.294 | 0.15 |
| RoBep | ESM-C | **0.379** | **0.340** | **0.234** | **0.268** | **0.330** | 0.35 |

**Figure 2.**
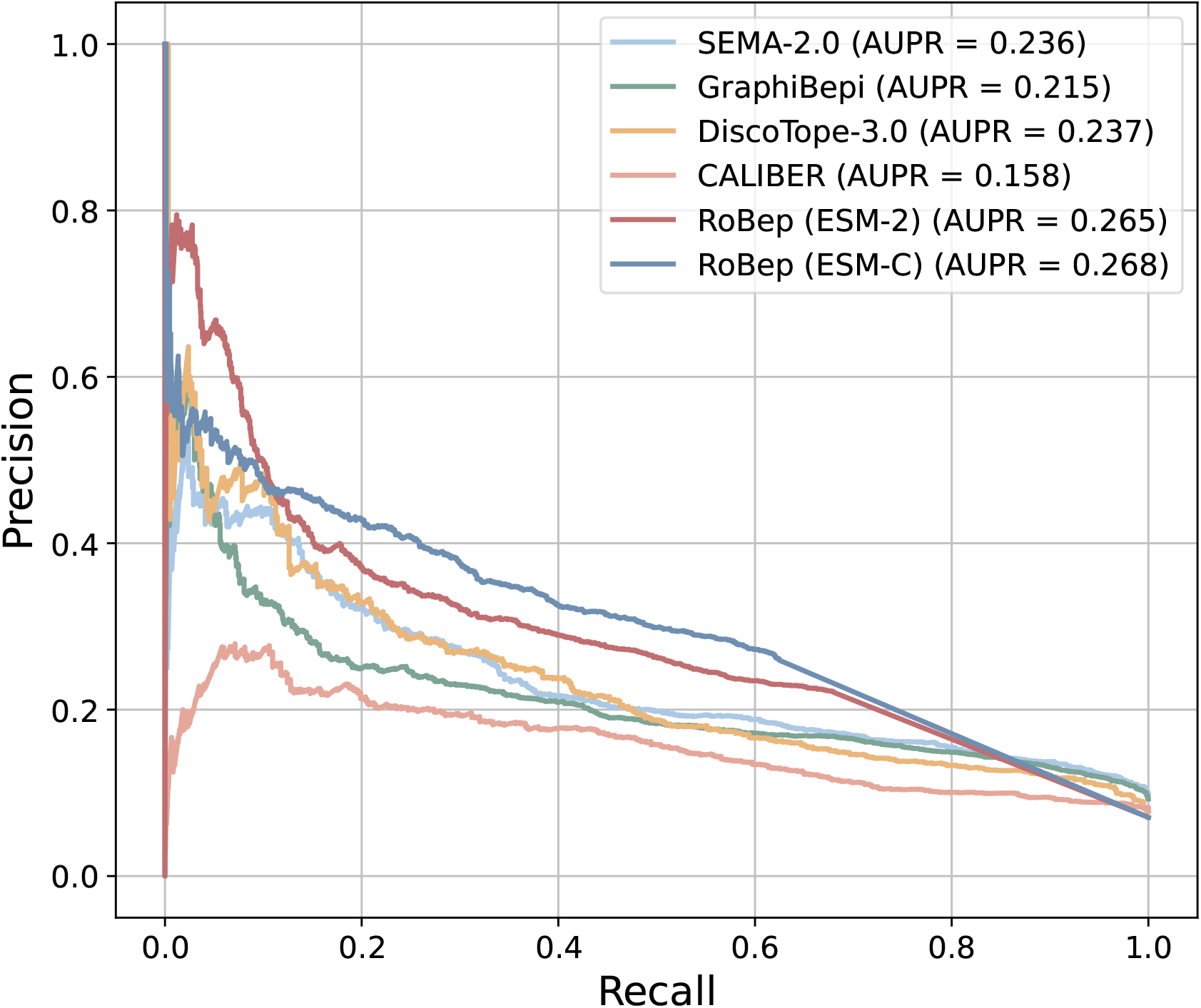
Precision–recall (AUPR) curves on the external test set. RoBep equipped with either ESM-2 (red) or ESM-C (indigo) consistently achieves higher precision across all recall levels.

Moreover, replacing the PLM with ESM-C, a model with more parameters and a more advanced architecture, led to further gains over the ESM-2-based version of RoBep: AUPRC increased by 0.9%, F1 score by 9.5%, AgIoU by 12.0%, and AUROC_0.1_ by 12.2%. These improvements are consistent with ESM-C’s enhanced sequence modeling capacity and further demonstrate that our model can effectively leverage high-quality embeddings to boost predictive performance.

Finally, the notably weaker performance of Seppa-3.0, which does not utilize any protein language model, underscores the importance of modern PLMs in capturing informative sequence or structure-derived representations.

### 3.2 Spatial Compactness of Predicted Epitope Residues

In this section, we evaluate the spatial compactness of epitopes predicted by each model, an important characteristic reflecting the biological plausibility and practical significance of predicted epitopes. To quantify compactness, we computed the clustered mean pairwise distance (cMPD) of predicted residues across each antigen in the independent test dataset, as detailed in Section 2.5. As shown in Figure 3, RoBep demonstrated significantly tighter spatial clustering compared to all baseline methods (*p <* 0.05, one-sided Dunnett’s test). Specifically, RoBep achieves an averaged cMPD of 18.49 Å with a standard deviation of 3.38 Å, closest to the distribution of true-epitope cMPD (13.62 Å ± 2.9 Å; red dashed line). In comparison, the averaged cMPDs for the baseline models are: CALIBER (21.91 Å), GraphiBepi (25.40 Å), SEMA-2.0 (25.91 Å), and DiscoTope-3.0 (28.02 Å).

**Figure 3.**
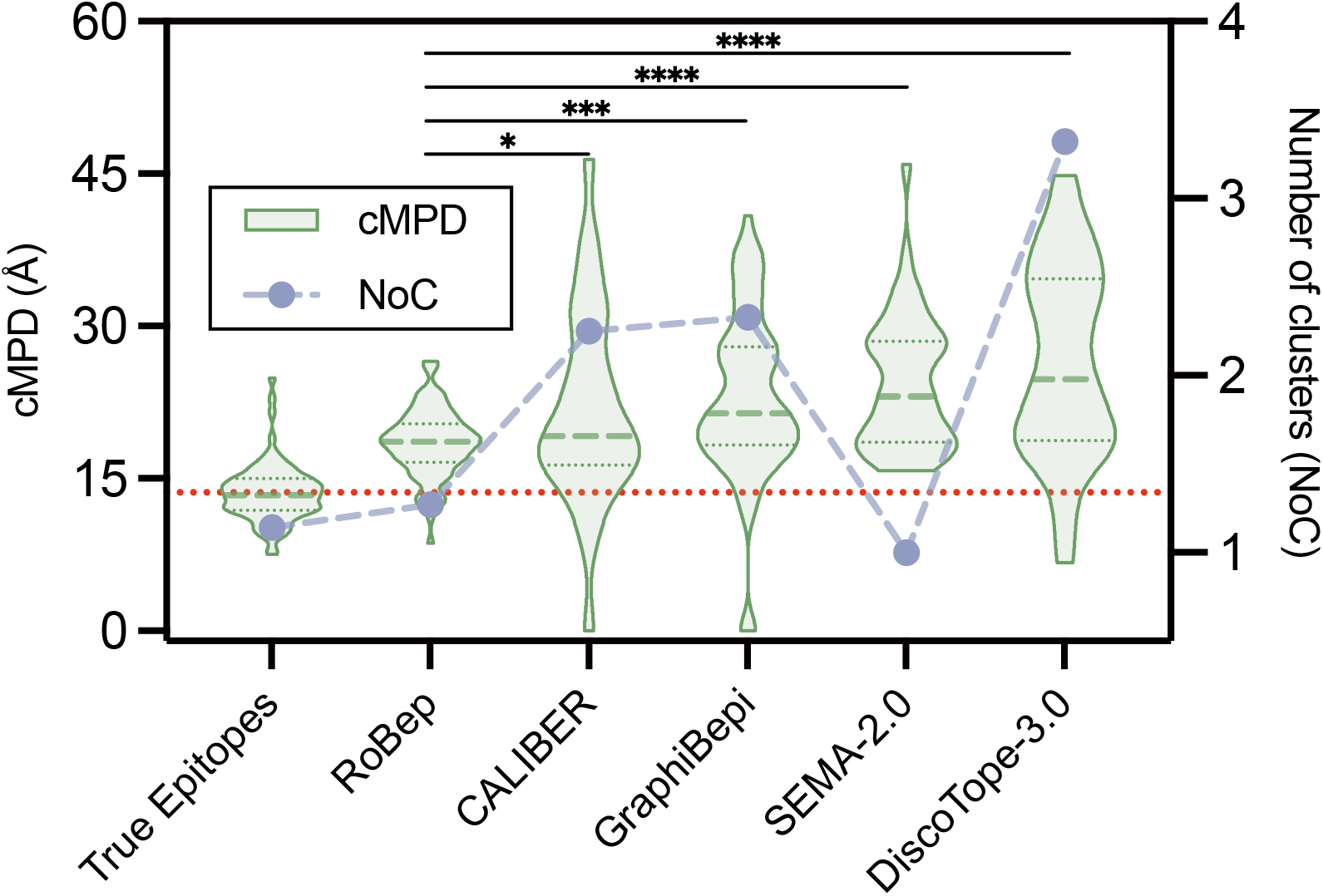
Comparison of cMPD and NoC among predicted epitopes and true epitopes on the independent test dataset. Lower cMPD values indicate tighter clustering. Asterisks denote statistical significance from one-sided Dunnett’s test against RoBep. The red dashed line indicates the mean cMPD of true epitopes across all test samples.

Beyond compactness, we also compare the number of clusters (NoC) implied by each model’s predictions to that of true epitopes. It can be found in Figure 3 that RoBep’s NoC is the closest to the ground truth, indicating preservation of epitope regional organization (avoiding both over-fragmentation into many small patches and over-merging into a single cluster). By contrast, SEMA-2.0 tends to produce one large connected cluster (NoC ≈ 1) with high cMPD, whereas other baselines tend to over-fragment epitopes. Together with the lower cMPD, these results demonstrate the effectiveness of our approach in producing spatially coherent predictions that align with biologically realistic epitope distributions.

### 3.3 Effect of Region Constraint and Other Components

To assess the contribution of region constraint and other modules in our architecture to BCE prediction, we conducted a series of ablation experiments. First, to evaluate the impact of region constraint, we removed the region enumeration and region ranking module to perform epitope prediction over the entire antigen, as in most existing methods. In this setting, the graph construction was changed from a region-based complete graph to a radius-based graph (18 Å cutoff) to maintain compatibility with EGNN. As visualized in Figure 4, removing the region constraint led to a substantial performance drop: F1 score fell from 0.3794 to 0.3247, MCC from 0.3398 to 0.2357, and AUPRC from 0.2676 to 0.2520. These decreases underscore that focusing predictions on a few high-likelihood surface patches both preserves biological consistency and significantly benefits BCE prediction.

**Figure 4.**
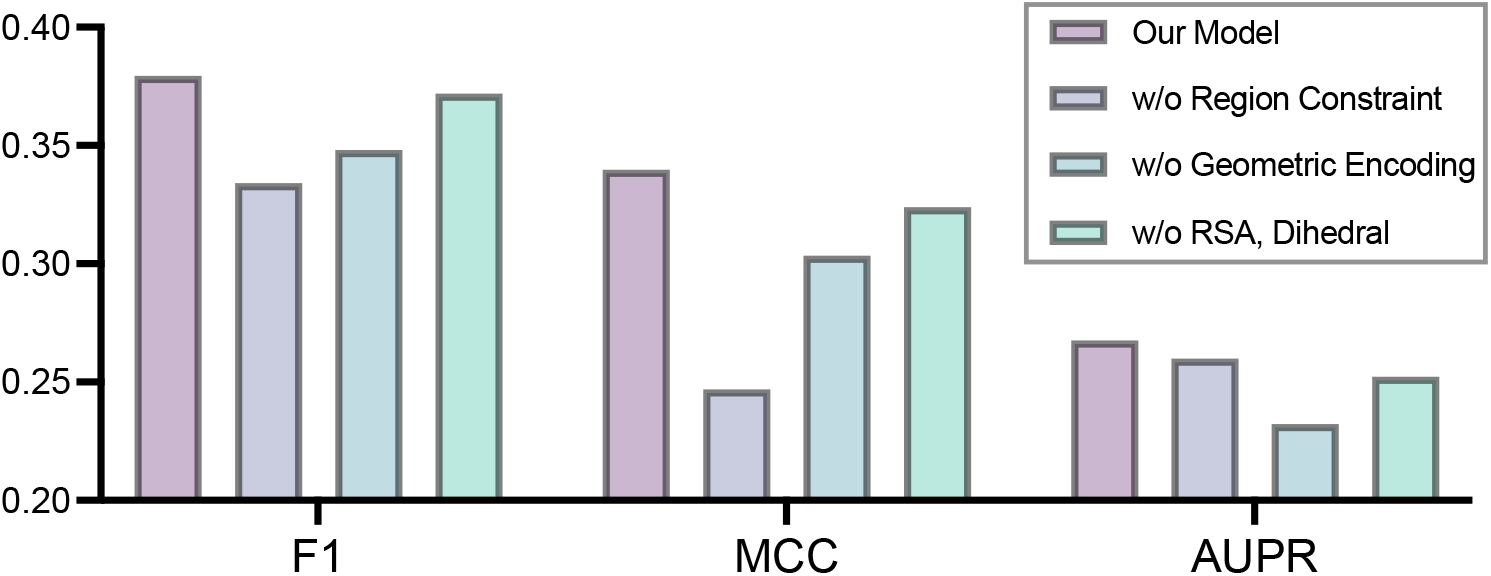
Performance comparison on the test dataset when removing Region Constraint or other individual components.

Next, omitting the Geometric Graph Encoding module caused notable declines in all metrics (F1 –8.2%, MCC –10.7%, AUPRC –13.2%), demonstrating its critical role in capturing spatial patterns. Finally, removing structural features, RSA and backbone torsion angles, resulted in a modest F1 drop of 2.0%, suggesting these descriptors provide complementary information, although some of their signal may already be captured by pretrained embeddings and geometric graph encoders.

### 3.4 Impact of Antigen Species and Length

Given the considerable diversity arising from antigen species and their variant lengths, we investigated how RoBep’s predictive performance varies across different antigen species and sequence lengths. Specifically, antigens were classified into three groups: (1) Human (autoantigens, tumor antigens, etc.), (2) Viral (all viral antigens), and (3) Other (including artificial proteins, Arrestin, etc.). We further divided them into five distinct length bins: 0–100, 100-200, 200-400, 400-700, and 700+ aa, for a more granular categorization based on antigen lengths. We computed mean AUPR per stratum as well as the number of non-redundant antigens in each stratum to reflect investigation density.

As shown in Figure 5, RoBep exhibits clear trends with respect to antigen length and species. First, although the 0–100 aa stratum contains fewer antigens than longer strata, it achieves the best performance (Viral: AUPR=0.569; Human: AUPR=0.491), and the mean AUPR decreases as length increases. This suggests that epitopes on shorter antigens are intrinsically easier to localize and may also reflect that representations learned on longer sequences transfer to shorter ones. Meanwhile, viral antigens consistently yielded higher AUPR values compared to human and other categories, even though human antigens are more numerous overall than viral (421 vs. 220 in our non-redundant set). It likely attributables to the fact that several viral antigens (e.g., influenza A [14] and coronaviruses [22]) contain relatively conserved epitope regions, whereas human antigens (including autoantigens [36] and tumor antigens [3]) are more heterogeneous across individuals and disease contexts. Notably, an intriguing performance rebound is observed for viral antigens in the 700+ aa bin (AUPR=0.491), likely driven by a higher concentration of well-studied viral antigens (e.g., spike-like proteins) that dominate this stratum. The overall trend indicates that shorter and viral antigen sequences may inherently contain epitope information that is easier to predict.

**Figure 5.**
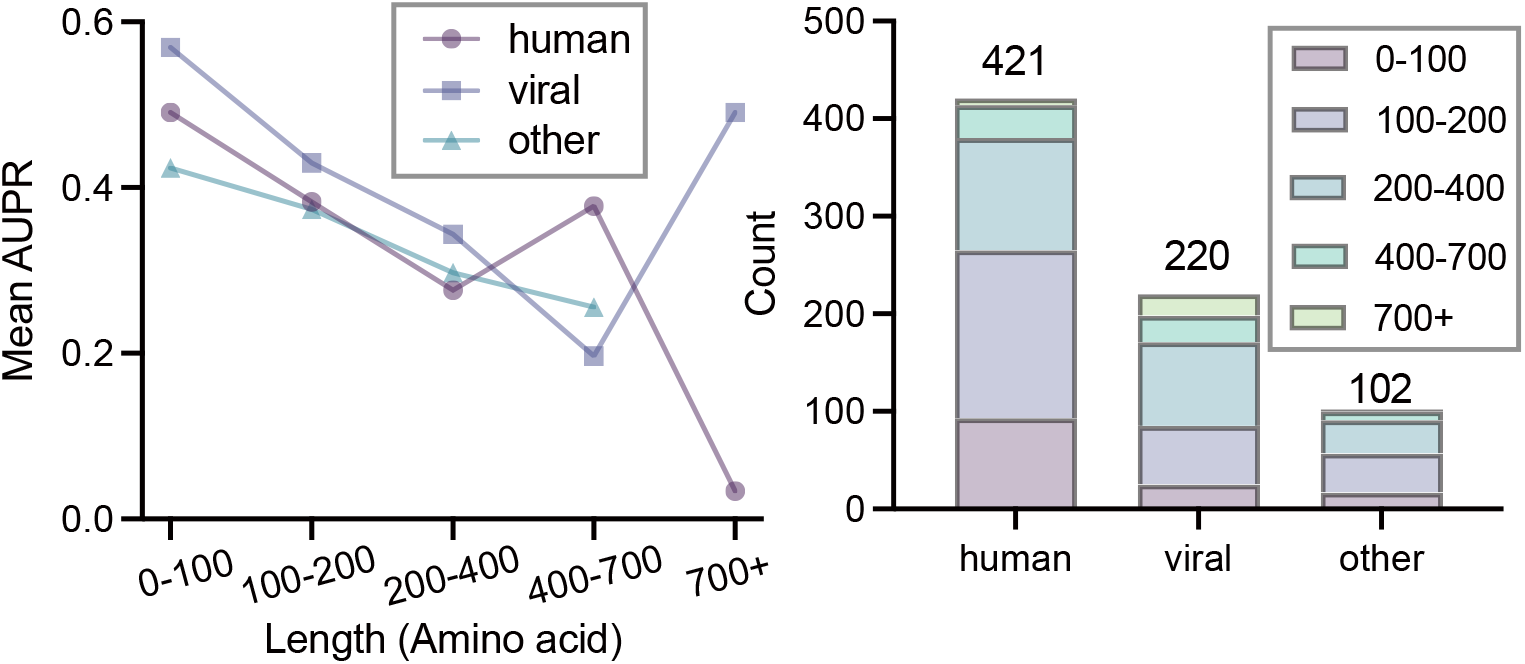
Mean AUPR of RoBep across species–length strata in the test set, with counts of non-redundant antigens per stratum.

### 3.5 Model Specificity on Antigenic Proteins

Identifying whether a given protein contains antigenic epitopes typically requires resource-intensive in vivo assays. Therefore, in silico methods capable of specifically recognizing antigenic proteins can greatly facilitate the initial screening of potential antigenic proteins. To evaluate RoBep’s specificity towards antigenic proteins, we examined its ability to distinguish known antigenic proteins from those relatively lacking antigenicity. Specifically, we conducted a binary classification on our antigenicity dataset (positives: antigens; negatives: viral nsps), using the protein-level antigenicity score defined in Section 2.4. The decision threshold (0.525) was selected on the training set (positives: antigens; negatives: their corresponding antibodies) to avoid data leakage and then fixed for evaluation on the antigenicity dataset introduced in Section 2.1.

As shown in Figure 6, RoBep assigned significantly lower antigenicity scores to most non-structural proteins, confirming its capacity to suppress false positives on weakly antigenic proteins. Our model also achieved strong discriminative performance, with AUPR = 0.9930, AUROC = 0.9905, F1 = 0.7500, and Recall = 0.6042. I It can be found that the model’s ability to identify antigenic proteins is weaker than that for non-antigenic proteins, yielding a lower recall, consistent with the intrinsic difficulty of generalizing across diverse antigen structures under limited data availability. Overall, these results confirm that RoBep effectively assigns lower antigenicity scores to proteins without antigenic properties, underscoring its practical value in specifically identifying genuine antigenic targets. More evaluation results on the antibody-antigen dataset are provided in Supplementary Materials S8.

**Figure 6.**
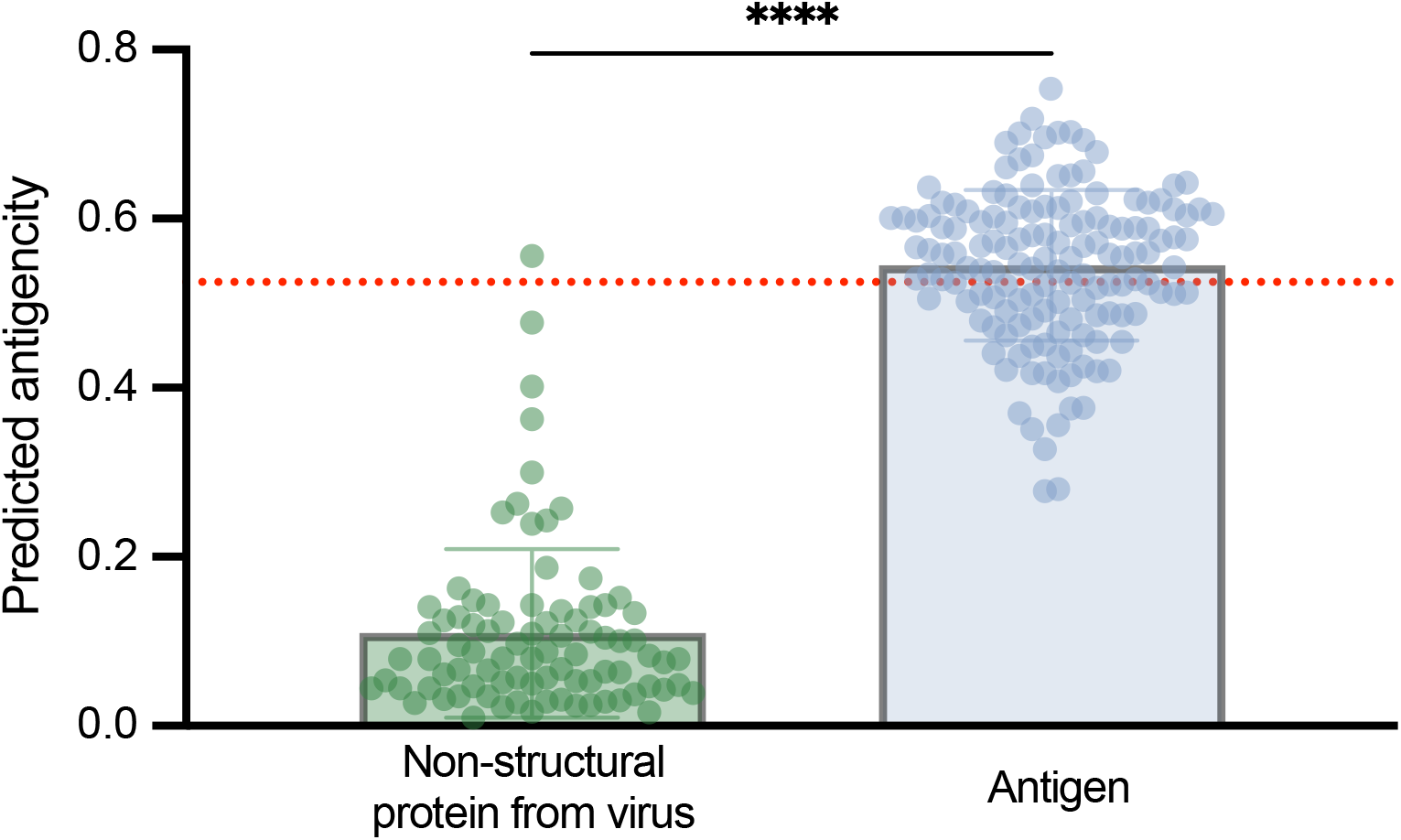
Predicted antigenicity of antigen and non-structural protein. The red dashed line indicates the threshold of 0.525.

### 3.6 Case Study

In this section, we present a representative case study on an antigen from the test set (PDB ID: 5i9q, Chain A) to further demonstrate the strength of our RoBep framework and the contribution of the region constraint mechanism. RoBep is compared with its variant without region constraint, as well as with four representative baselines: GraphBepi, SEMA-2.0, DiscoTope-3.0, and CALIBER.

As shown in Figure 7, RoBep achieves the highest performance across all metrics, with an F1 score of 0.88, AUPR of 0.75, and precision of 0.86. Notably, even without the region constraint mechanism, our model maintains competitive performance, confirming the robustness of its underlying architecture. However, this unconstrained variant exhibits a notable increase in false positives (red), underscoring the value of the region constraint in suppressing noisy predictions and enforcing spatial focus.

**Figure 7.**
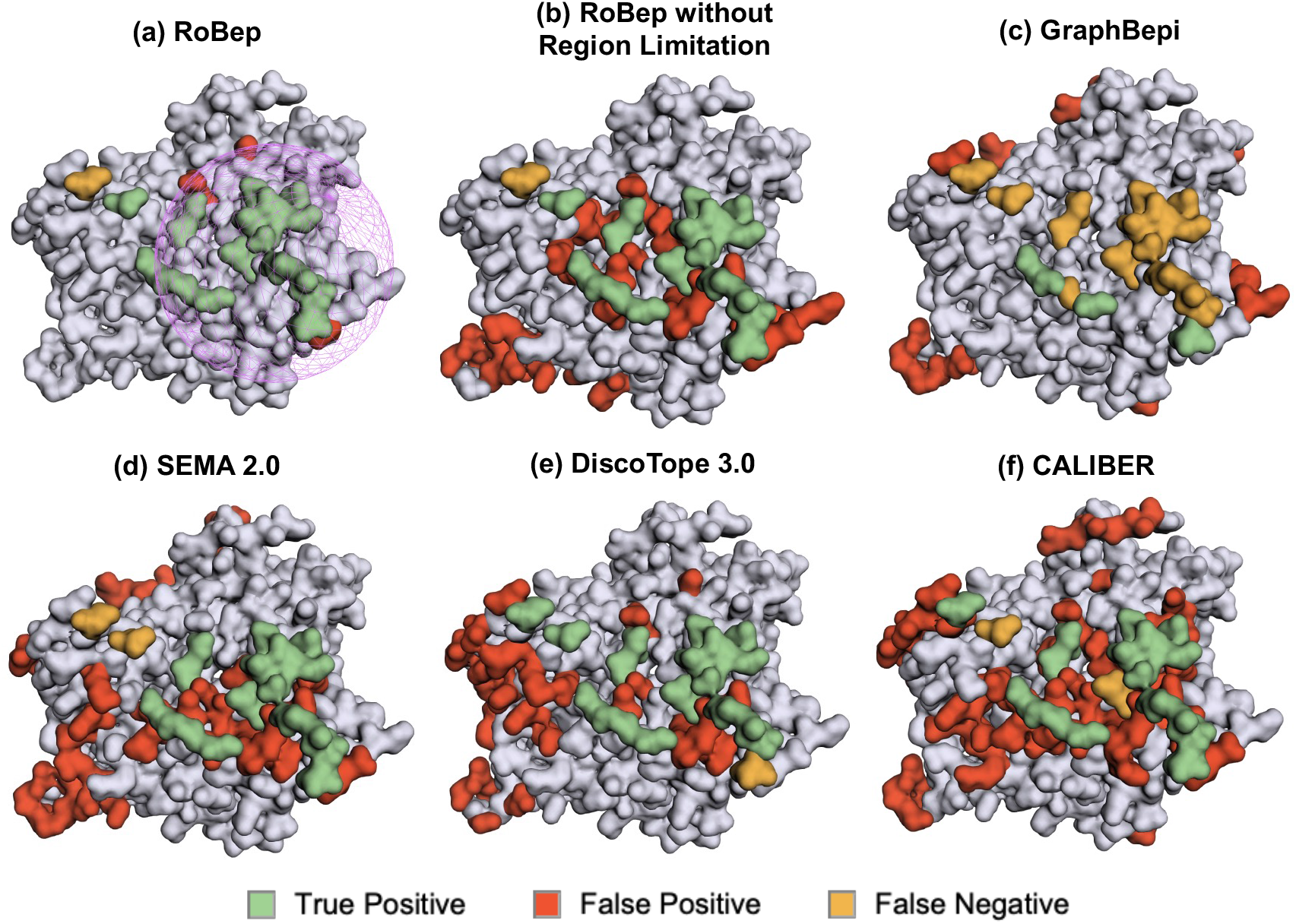
Visual comparison of predicted B-cell epitopes on antigen 5I9Q (chain A) by six methods: (a) RoBep, (b) RoBep without Region Constraint, (c) GraphBepi, (d) SEMA-2.0, (e) DiscoTope-3.0, and (f) CALIBER. The purple sphere indicates the region with the highest predicted likelihood of being an antibody–antigen interface, as identified by RoBep.

In comparison, baseline methods such as SEMA-2.0, DiscoTope-3.0, and CALIBER generate predictions that are more scattered across the antigen surface, despite identifying a comparable number of true epitope residues. This results in substantially lower precision (0.38, 0.53, and 0.36, respectively) and weaker spatial coherence. By contrast, RoBep produces highly spatially clustered predictions, closely aligning with how B-cell epitopes are typically present in nature, as compact regions. This spatial compactness not only improves biological plausibility but also provides more actionable guidance for downstream tasks such as vaccine design and antibody development. More examples with different performance can be found in the Supplementary S9.

## 4 Discussion

In this study, we introduced RoBep, a region-constrained framework for B-cell epitope prediction. By explicitly modeling the spatial clustering of epitopes through a region constraint mechanism, RoBep generates biologically coherent predictions and effectively avoids the overly scattered outputs commonly observed in existing residue-level models. The framework integrates ESM-C and equivariant graph neural networks (EGNN) to jointly capture informative sequence and structural representations. In addition to providing residue-level epitope probabilities, RoBep also predicts high-likelihood antibody–antigen binding regions, making it a practical and interpretable tool for structure-guided vaccine design and antibody discovery. Notably, RoBep demonstrates a degree of specificity on antigenic proteins and achieves consistently strong performance in benchmark comparison. Note that RoBep can also predict linear BCEs since linear B-cell epitope residues are also clustered on an antigen surface region.

Despite its advantages, RoBep has certain limitations. First, its region enumeration strategy is based on a fixed-radius spherical scanning approach, which may not well adapt to antigens of varying size, shape, or domain architecture. Future work may explore more adaptive or data-driven region selection strategies to improve coverage and generalizability. Second, while the region constraint enforces spatial coherence, it introduces a high-risk high-reward trade-off: if all top-ranked regions miss the true epitope location, the model has limited capacity to recover correct predictions. This limitation suggests the possibilities of incorporating hierarchical or soft region constraints to improve robustness without sacrificing spatial focus, allowing the model to make more flexible and resilient predictions.

In summary, RoBep is the first biologically grounded, structure-aware framework to incorporate explicit spatial constraints into B-cell epitope prediction. By combining advanced deep learning models with a region constraint mechanism, RoBep not only achieves accurate and biologically plausible residue-level outputs but also introduces region-level BCE identification for the first time. Beyond epitope prediction, we believe the region constraint strategy may serve as a general design principle for other structure-based localization tasks where spatial coherence and interpretability are essential.

## Supplementary Data

Supplementary Data are available at *Bioinformatics* online. Conflict of interest: None declared.

## Footnotes

1 conceptually related to the classical rolling-sphere view of solvent exposure [26]

2 Residues with relative solvent accessibility (RSA) ≥ 20%.

