## Supplementary Materials for "RoBep: A Region-Oriented Deep Learning Model for B-Cell Epitope Prediction"


#### S1. External Information of Dataset

##### S1.1 Statistics of the Antibody-Antigen (Ab-Ag) Complexes Uploaded to the PDB Database Across Different Species

Figure 1 summarizes the number of antigen–antibody complex structures deposited in the PDB each year since 1989, covering all antigens with sequence lengths ranging from 12 to 2046 residues and resolution better than 4.0 Å, across different species (3,454 in total).

##### S1.2 Construction of Non-Structural Proteins Dataset

All 85 non-structural proteins from viruses used to assess RoBep's ability to distinguish antigenic proteins from weakly antigenic NSPs were curated from the Protein Data Bank (PDB) Berman et al. [2000] using the keywords '`virus AND (enzyme OR protease OR polymerase OR helicase)`', as most viral enzymes are non-structural, particularly during early stages of viral formation Murphy et al. [1999]. Each candidate was manually verified to exclude structural proteins and proteins may be antigenic. The full list is available in our GitHub repository: https://github.com/YitaoXU/RoBep.

#### S2. Feature Construction Details

##### S2.1 Node Features

For each residue $i$ in a candidate region, we extract three types of features:

**(1) ESM-C embeddings.** We use ESM Cambrian 6b [Hayes et al., 2025], a protein language model trained via masked language modeling, to produce a 2560-dimensional embedding $\mathbf{x}_i^{\text{ESM}} \in$

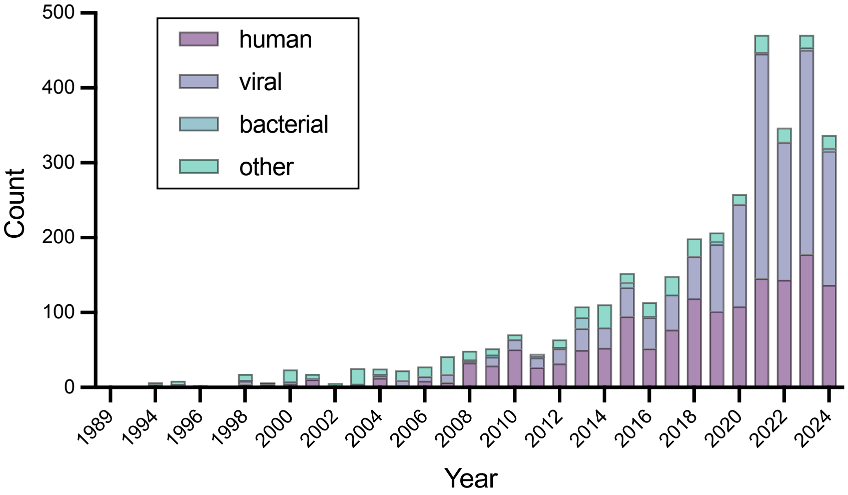


Figure 1: Number of Ab-Ag complexes uploaded to the PDB database across different species

$\mathbb{R}^{2560}$, which is then projected to the model's hidden dimension via a learnable MLP:

$$\mathbf{h}_i^{\mathrm{ESM}} = \phi_{\mathrm{ESM}}(\mathbf{x}_i^{\mathrm{ESM}}) \in \mathbb{R}^{d_{\mathrm{ESM}}}$$

**(2) Relative solvent accessibility (RSA).** RSA is computed using the Shrake–Rupley algorithm [Shrake and Rupley, 1973] implemented in Biotite [Kunzmann and Hamacher, 2018], and encoded as a scalar:

$$\mathbf{h}_i^{\mathrm{RSA}} \in \mathbb{R}^1$$

**(3) Dihedral angles.** Backbone torsion angles $(\phi_i, \psi_i, \omega_i)$ are computed from atomic coordinates and encoded via sine and cosine functions:

$$\mathbf{h}_i^{\mathrm{dihedral}} = [\sin\phi_i, \cos\phi_i, \sin\psi_i, \cos\psi_i, \sin\omega_i, \cos\omega_i] \in \mathbb{R}^6$$

All components are concatenated to form the initial node embedding:

$$\mathbf{h}_i^{(0)} = [\mathbf{h}_i^{\mathrm{ESM}} \parallel \mathbf{h}_i^{\mathrm{RSA}} \parallel \mathbf{h}_i^{\mathrm{dihedral}}]$$

with dimension $d_n = d_{\mathrm{ESM}} + 1 + 6$ and $d_{\mathrm{ESM}} = 512$ in final model.

##### S2.2 Edge Features

Each edge $(i, j)$ connects two residues in the same region and includes:

**(1) Radial distance.** The Euclidean distance between $C_\alpha$ atoms is expanded using 16 Gaussian radial basis functions:

$$\mathbf{a}_{ij}^{\mathrm{RBF}} \in \mathbb{R}^{16}$$

**(2) Sequence offset.** The residue index difference $j - i$ is encoded as a sinusoidal positional embedding [Vaswani et al., 2017]:

$$\mathbf{a}_{ij}^{\sin} \in \mathbb{R}^{16}$$

The final edge embedding is:

$$\mathbf{a}_{ij} = [\mathbf{a}_{ij}^{\mathrm{RBF}} \parallel \mathbf{a}_{ij}^{\sin}] \in \mathbb{R}^{32}$$

#### S3 Selection of $k$ in Region Scoring and Ranking

To determine the optimal number of top-ranked regions ($k$) for residue-level prediction, we conducted a grid search over $k$ from 1 to 20 using 3-fold cross-validation on the training set. The data was evenly partitioned into three folds; in each round, two folds were used for training and the remaining one for validation. For each $k$ value, we trained models on the training folds and evaluated them on the corresponding validation fold. The validation AUPRC was averaged across the three folds, and we found that $k = 7$ yielded the highest overall score. The table below summarizes the validation results for different $k$ values:

| $k$ | AUPRC |
|---|---|
| 1 | 0.2947 |
| 2 | 0.3026 |
| 3 | 0.3098 |
| 4 | 0.3239 |
| 5 | 0.3285 |
| 6 | 0.3326 |
| **7** | **0.3356*** |
| 8 | 0.3314 |
| 9 | 0.3277 |
| 10 | 0.3275 |
| 11 | 0.3287 |
| 12 | 0.3300 |
| 13 | 0.3318 |
| 14 | 0.3279 |
| 15 | 0.3295 |
| 16 | 0.3290 |
| 17 | 0.3281 |
| 18 | 0.3275 |
| 19 | 0.3248 |
| 20 | 0.3240 |

Table 1: Validation AUPRC for different numbers of top-ranked regions ($k$) on the held-out 20% validation set. * indicates the highest AUPRC.

#### S4. GradNorm for Dynamic Loss Balancing

To dynamically balance region- and residue-level supervision during training, we adopt GradNorm [Chen et al., 2018], a gradient normalization strategy designed for multi-task learning.

GradNorm dynamically adjusts the loss weights to ensure that different tasks train at a similar rate. Let $\mathcal{L}_1$ and $\mathcal{L}_2$ denote two task-specific losses, and $w_1$, $w_2$ their corresponding learnable weights. The gradients of the weighted losses with respect to the shared parameters $\mathbf{W}$ (e.g., MLP for ESM-C embeddings, EGNN encoder) are computed as:

$$G_i(t) = \left\| \nabla_{\mathbf{W}} \left( w_i(t) \cdot \mathcal{L}_i(t) \right) \right\|_2 , \quad i \in \{1, 2\}$$

, where $G_i(t)$ is the $L_2$ norm of the gradient for task $i$ at training step $t$.

To guide weight updates, GradNorm defines a target gradient norm $\hat{G}_i(t)$ for each task based on its relative training progress:

$$\hat{G}_i(t) = \bar{G}(t) \cdot \left( \frac{\tilde{\mathcal{L}}_i(t)}{\bar{\mathcal{L}}(t)} \right)^{\alpha}$$

, where:$\tilde{\mathcal{L}}_i(t) = \mathcal{L}_i(t)/\mathcal{L}_i(0)$ is the normalized loss value of task $i$; $\bar{G}(t) = \frac{1}{2}(G_1(t) + G_2(t))$ is the average gradient norm across tasks; $\bar{\mathcal{L}}(t) = \frac{1}{2}(\tilde{\mathcal{L}}_1(t) + \tilde{\mathcal{L}}_2(t))$ is the average normalized loss; $\alpha$ is a hyperparameter controlling how strongly to penalize imbalance.

GradNorm updates $w_1, w_2$ by minimizing the following auxiliary loss:

$$\mathcal{L}_{\text{grad}} = \sum_{i=1}^{2} \left| G_i(t) - \hat{G}_i(t) \right|$$

. This encourages the gradient norms of all tasks to stay close to their targets. After updating, the weights are renormalized to keep their sum constant.

**Implementation details.** In RoBep, we apply GradNorm to balance the region-level loss $\mathcal{L}_{\text{region}}$ and the residue-level loss $\mathcal{L}_{\text{node}}$, using gradients computed with respect to the shared parameters. We initialize the weights ratio $\frac{w_{\text{node}}}{w_{\text{region}}} = 0.5$ to prioritize region-level supervision at early training. GradNorm updates are performed every 10 training steps.

#### S5. Evaluation Metrics

Let $\text{TP}, \text{TN}, \text{FP}, \text{FN}$ denote the number of true positives, true negatives, false positives, and false negatives in residue-level epitope prediction. Let $\mathcal{E} = \{\mathbf{r}_1, \ldots, \mathbf{r}_n\} \subset \mathbb{R}^3$ be the set of $\text{C}_\alpha$ coordinates of the $n$ predicted epitope residues.

##### F1 score

$$\text{F1} \;=\; \frac{2\,\text{TP}}{2\,\text{TP} + \text{FP} + \text{FN}}.$$

The F1 score balances precision and recall, offering a robust evaluation for imbalanced datasets.

##### Matthews correlation coefficient (MCC)

$$\text{MCC} \;=\; \frac{\text{TP} \times \text{TN} - \text{FP} \times \text{FN}}{\sqrt{(\text{TP} + \text{FP})(\text{TP} + \text{FN})(\text{TN} + \text{FP})(\text{TN} + \text{FN})}}.$$

MCC accounts for all elements in the confusion matrix and remains reliable even under class imbalance.

##### Antigen-level intersection over union (AgIoU)

$$\text{AgIoU} \;=\; \frac{\text{TP}}{\text{TP} + \text{FP} + \text{FN}}.$$

AgIoU [You et al., 2025], which is the same as Jaccard Index, evaluates the overlap between predicted and true epitope residues at the antigen level by calculating the ratio of the number of correctly predicted epitope residues to the total number of residues that are either labeled as epitope in the ground truth or prediction.

**Area under the precision-recall curve (AUPR)**

$$\mathrm{AUPR} = \int_0^1 \mathrm{Precision}(r) \cdot \frac{d}{dr}\mathrm{Recall}(r)\, dr$$

AUPR evaluates the trade-off between precision and recall across classification thresholds, and is particularly informative in imbalanced settings where positive samples are rare.

**Area under the ROC curve (AUROC)**

$$\mathrm{AUROC} = \int_0^1 \mathrm{TP}(f) \cdot \frac{d}{df}\mathrm{FP}(f)\, df$$

AUROC measures the model's ability to rank positive instances above negative ones and reflects the probability that a randomly chosen positive is ranked higher than a randomly chosen negative.

**Area under the ROC curve in low-FPR region ($\mathrm{AUROC}_{0.1}$)**

$$\mathrm{AUROC}_{0.1} = \int_0^{0.1} \mathrm{TP}(f) \cdot \frac{d}{df}\mathrm{FP}(f)\, df$$

$\mathrm{AUROC}_{0.1}$ focuses on the low false-positive rate region ($\mathrm{FPR} \in [0, 0.1]$), emphasizing early retrieval performance that is particularly relevant for highly imbalanced residue-level prediction tasks.

**Clustered Mean pairwise distance (cMPD) and Number of Clusters (NoC)** This is a metric modified from Colavin et al. [2022] that provides a fair and accurate assessment of residues compactness, including multi-region cases. For each antigen $i$, we compute the cMPD of predicted epitopes in the following three steps:

1. Infer the ground-truth number of epitope regions.
   (a) Build an undirected graph $G_i = (V, E)$ with nodes $V = E_i^{\mathrm{true}}$ (ground-truth epitope residues).
   (b) Connect residues $r, s \in V$ with an edge if the minimum distance between any heavy atom of $r$ and any heavy atom of $s$ is $\leq 10$ Å. The selection of 10 Å cutoff follows the common practice that residues within $\leq 10$ Å are considered proximal in proteins Jing and Xu [2021], Gligorijević et al. [2021].
   (c) Run breadth-first search to extract the connected components $\{C_{i,1}^{\mathrm{true}}, \ldots, C_{i,k_i}^{\mathrm{true}}\}$ of $G_i$ and set $k_i$ to the number of clusters (NoC).
2. Cluster the predicted epitope residues.
   (a) Apply $k$-means clustering to the predicted epitope residues $E_i^{\mathrm{pred}}$ using their C$\alpha$ coordinates, with $k = k_i$.
   (b) Denote the resulting clusters by $\{C_{i,1}^{\mathrm{pred}}, \ldots, C_{i,k_i}^{\mathrm{pred}}\}$.
3. Compute MPD within clusters and aggregate with pair-wise weighting.

(a) For each cluster $C_{i,j}^{\text{pred}}$, let $n_j = |C_{i,j}^{\text{pred}}|$ and $\{\mathbf{c}_1, \ldots, \mathbf{c}_{n_j}\}$ be the C$\alpha$ coordinates. Define the mean pairwise distance as

$$\text{MPD}_{i,j} = \begin{cases} 0, & n_j < 2, \\ \dfrac{2}{n_j(n_j-1)} \displaystyle\sum_{1 \le p < q \le n_j} \|\mathbf{c}_p - \mathbf{c}_q\|_2, & n_j \ge 2. \end{cases}$$

(b) The Clustered MPD for antigen $i$ is computed as a weighted average, where each cluster is weighted by its number of residue pairs $\binom{n_j}{2}$:

$$\text{cMPD}_i = \frac{\displaystyle\sum_{j=1}^{k_i} \binom{n_j}{2} \cdot \text{MPD}_{i,j}}{\displaystyle\sum_{j=1}^{k_i} \binom{n_j}{2}} = \frac{\displaystyle\sum_{j=1}^{k_i} \frac{n_j(n_j-1)}{2} \cdot \text{MPD}_{i,j}}{\displaystyle\sum_{j=1}^{k_i} \frac{n_j(n_j-1)}{2}}.$$

This weighting scheme ensures that larger clusters (with more residue pairs) contribute more to the overall metric.

This design ensures that when the ground truth contains multiple regions, correctly predicting two or more regions on opposite sides is not penalized. Conversely, when the ground truth has a single region.Colavin et al. [2022]. Smaller values indicate tighter, more coherent clustering on the antigen surface.

#### S6. Training Implementation Details

To identify the best hyperparameter setting, we performed 5-fold cross-validation on the training set. Specifically, the training data was evenly split into three subsets. In each fold, two subsets were used for training and the remaining one for validation. We repeated this process three times, each time using a different subset as the validation set, and selected the hyperparameter configuration that achieved the highest average AUPR across the three folds through grid search. The hyperparameter search space is summarized in the Table 2.

| **Hyperparameter** | **Search Space** |
|---|---|
| ESM-C projected dimension | {374, **512**, 748, 1024} |
| EGNN hidden layers | {[512], [512,512], [512,512,512],<br>[512,256], [512,256,256], [**256,256**]} |
| Number of node classifier layers | {1, **2**, 3, 4} |
| Consistency loss weight | {0, 0.1, 0.2, **0.3**, 0.4} |
| Node:Region loss weight ratio | {0.1, **0.5**, 1.0, 2.0, 5.0} |
| Region type | {**Weighted MSE**, BCE, Combined loss (weighted MSE + BCE).} |
| Learning rate | {5e-4, 2e-4, **5e-5**, 1e-6} |

Table 2: Hyperparameter search space for model selection, with the selected parameters in bold.

| Metric | RoBep (Weighted MSE) | RoBep (BCE) | RoBep (Combined loss) |
|---|---|---|---|
| AUPRC | **0.2675** | 0.2526 | 0.2655 |
| AUROC | 0.7562 | **0.7692** | 0.7539 |
| $AUROC_{0.1}$ | **0.3303** | 0.2813 | 0.3302 |
| Accuracy | **0.8664** | 0.8051 | 0.8558 |
| Precision | **0.2823** | 0.2173 | 0.2696 |
| Recall | 0.5783 | **0.6766** | 0.6095 |
| F1-Score | **0.3794** | 0.3289 | 0.3738 |
| AgIoU | **0.2341** | 0.1968 | 0.2299 |
| MCC | **0.3398** | 0.3040 | 0.3385 |

Table 3: Performance comparison between RoBep trained with different region losses: MSE-only loss, BCE-only loss, and Combined (MSE + BCE) loss.

| | AUPR | F1 | MCC |
|---|---|---|---|
| RoBep | 0.525 | 0.524 | 0.203 |
| Random Selection | 0.331 (0.333) | 0.325 (0.333) | -0.014 (0.000) |

Table 4: Comparison of model specificity on antigenic proteins with random selection. The values in the parentheses are the mathematical expectations for these metrics.

### S7. Extra Evaluation Results on Different Choices of Loss

We labeled a region as positive if $y_{\text{region}_j} > 0.5$, and compared three variants of the region-level loss: (a) weighted MSE (our default), (b) BCE on the binarized target, and (c) a combined loss (weighted MSE + BCE). It can be found in Table 3 that weighted MSE yielded the best overall performance across most metrics, with the combined loss performing similarly, while BCE had higher recall and full AUROC at the cost of precision-oriented metrics.

### S8. Extra Evaluation Results of Model Specificity on Antigenic Proteins

Antigens from the test set served as positive examples, while corresponding antibody heavy and light chains were designated as negative examples in this evaluation. Given that antibodies inherently lack antigenicity. This yields a positive-to-negative ratio of 1:2.Since there are no existing methods available for this specific evaluation scenario, RoBep was benchmarked against a random selection baseline matching the dataset's class proportions. As a result, our model achieved an AUPR of 0.525, an F1 of 0.524, and an MCC of 0.203, significantly exceeding the random baseline (AUPR: 0.333, F1: 0.333, MCC: 0).

### S9. Case Study on Extra Antigen

To broaden performance coverage, we further consider two more examples. The first one is a randomly sampled extra Ab–Ag complex from the August 2025 PDB release (PDB: 9IRC Zhu et al. [2025]), whose antigen shares $< 70\%$ sequence identity to any training antigen (CD-HIT-2D). On this example, RoBep achieves AUPR:0.542, $AUROC_{0.1}$:0.675, F1:0.448, and AgIoU:0.289,

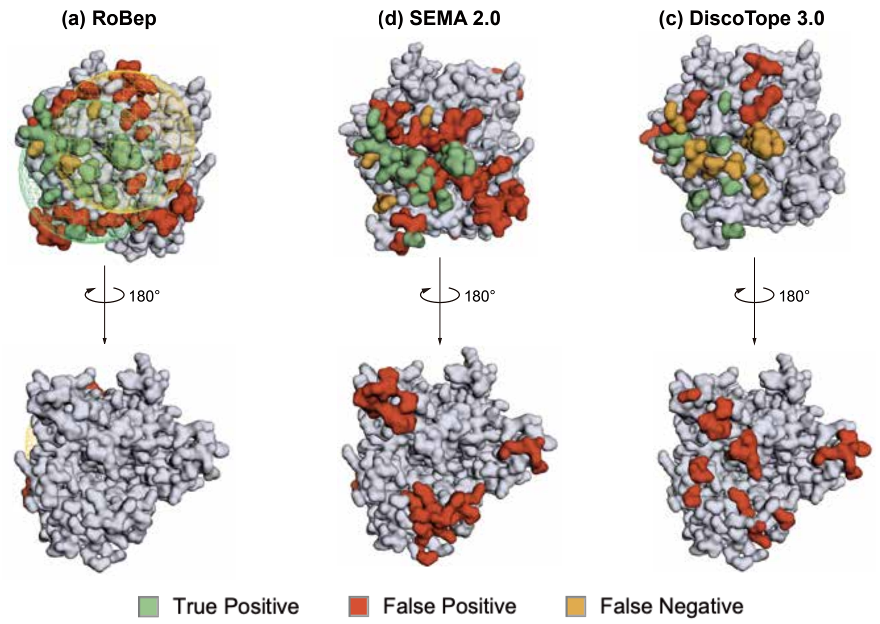


Figure 2: Visual comparison of predicted B-cell epitopes on antigen 9irc (chain A) by three methods: (a) RoBep, (b) SEMA-2.0, and (c) DiscoTope-3.0 The yellow and green spheres indicate the region with the highest two predicted likelihood of being an antibody–antigen interface, as identified by RoBep.

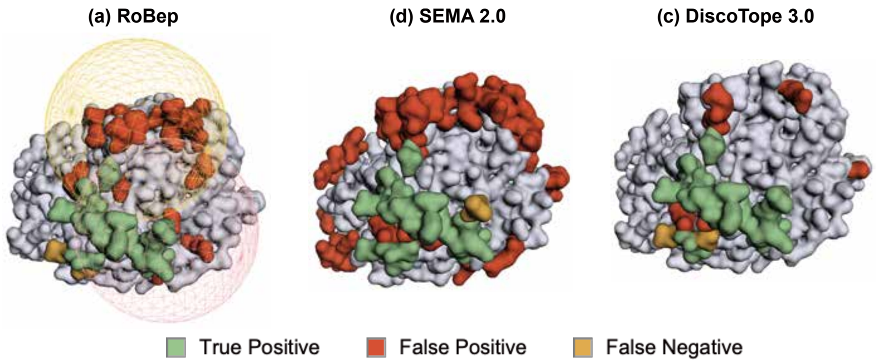


Figure 3: Visual comparison of predicted B-cell epitopes on antigen 9irc (chain A) by three methods: (a) RoBep, (b) SEMA-2.0, and (c) DiscoTope-3.0 The yellow and red spheres indicate the region with the highest two predicted likelihood of being an antibody–antigen interface, as identified by RoBep.

outperforming DiscoTope-3.0 (AUPR:0.279, $\text{AUROC}_{0.1}$:0.501, F1:0.100, AgIoU:0.053) and SEMA-3.0 (AUPR:0.355, $\text{AUROC}_{0.1}$:0.524, F1:0.364, AgIoU:0.222). As shown in Figure 2, RoBep's region-constraint design yields compact, localized predictions that avoid surface-wide dispersion, even though the false positive rate in selected region is not that low.

Second one is 3T2N(A), a serine protease hepsin picked from our test set. Our model attains AUPR=0.480, AUROC0.1=0.329, F1=0.596, and AgIoU=0.424, a little lower than Discotope (AUPR=0.751, AUROC0.1=0.339, F1=0.667, AgIoU=0.5). This gap is attributable to our current fixed sphere count/radius, which are relatively large for this target and increase false positives (Fig. 3). As noted in the Discussion, we will adopt adaptive/data-driven region selection (e.g., dynamically varying sphere radius or alternative shapes) to improve coverage and generalizability. RoBep nonetheless still reduces false negatives via its region-constraint mechanism to some extent compared with SEMA-3.0.
